# Neural Signatures of Emotion Regulation

**DOI:** 10.1101/2023.06.12.544668

**Authors:** Jared Rieck, Julia Wrobel, Antonio R. Porras, Kateri McRae, Joshua Gowin

## Abstract

Emotional experience is central to a fulfilling life. Although exposure to negative experiences is inevitable, an individual’s emotion regulation response may buffer against psychopathology. Identification of neural activation patterns associated with emotion regulation via an fMRI task is a promising and non-invasive means of furthering our understanding of the how the brain engages with negative experiences. Prior work has applied multivariate pattern analysis to identify signatures of response to negative emotion-inducing images; we adapt these techniques to establish novel neural signatures associated with conscious efforts to modulate emotional response. We model voxel-level activation via LASSO principal components regression and linear discriminant analysis to predict if a subject was engaged in emotion regulation and to identify brain regions which define this emotion regulation signature. We train our models using 82 participants and evaluate them on a holdout sample of 40 participants, demonstrating an accuracy up to 82.5% across three classes. Our results suggest that emotional regulation produces a unique signature that is differentiable from passive viewing of negative imagery.

## Introduction

Emotion regulation—the ability to change our experience or expression of emotions—is a fundamental human process and has a profound impact on mental health^1^. For example, the strategies that individuals tend to use for emotion regulation can reduce the likelihood of psychopathology when effective, or predispose individuals to developing depression, anxiety, or substance use disorders when they go awry^2,3^. One common emotion regulation strategy, cognitive reappraisal, involves thinking differently about a potentially emotion-inducing situation to alter its emotional impact^4^. Strengthening this skill is the goal of many empirically supported therapeutic interventions, and it forms the basis of cognitive behavioral therapy^5,6^.

While emotion regulation is a clinical target that has been studied experimentally, there re- mains a need to better understand its biological basis. Numerous neuroimaging studies in humans have explored the circuitry that contributes to emotional response to negative imagery^7,8^and circuitry involved in modifying a natural response using cognitive reappraisal^9,10,11^. Empirical work has supported theoretical neural models of emotion^12^, showing that regions such asthe amygdala^13^, insula^14^, and striatum^15^ encode salience and negative emotional reactions. The activity of these regions can then be modified by higher-order processes initiated in the anterior cingulate^16^, dorsomedial and dorsolateral prefrontal cortex, and parietal cortices^10^. Negative emotional response may be elevated in adults with mental health problems such as depression^17^, and the higher order influence of emotional regulation may be diminished in adults with psychopathologies^18^. Therefore, understanding the biological bases of both negative emotional response and our ability to control it using emotion regulation is relevant to understanding mental health.

Despite the progress in understanding emotion regulation, much remains unclear. Like many broad neural processes, such as craving^19^ and pain^20^, the neural representation of emotion spans multiple regions and cannot be summarized by the activation of a single structure. A neural signature for picture-induced negative emotion shows that many brain regions contribute to the expression of negative emotion and in aggregate they are highly sensitive and specific to reflecting negative emotion^21^. While Chang et al. establishes a neural signature of negative emotional response, our goal is to identify a neural signature that indicates when a person is actively engaging in cognitive processes to modulate their negative emotional response to pictures, which we refer to as reappraisal.

A signature of reappraisal could serve as a biomarker for diagnosis of mental health disorders. It could also serve as a therapeutic target for interventions and one metric of treatment success. Most reappraisal tasks rely upon self-reported negative emotion as their primary measurement, but self-reported emotion is vulnerable to lack of self-awareness of one’s emotions, deliberate deception, and lack of shared linguistic representations of emotion. Therefore, access to a neuroimaging measure as a critical methodological tool that is not subject to the same weaknesses represents a potential methodological advance^22,23^. Yet, to date such a signature has not been developed.

Here, we develop the first signature of reappraisal in data from 82 healthy young adults. In developing this signature, we compare the results of two machine learning models: the commonly employed LASSO principal component regression (LASSO PCR) and linear discriminant analysis (LDA). Specifically, we train the models to classify images into three categories defined by an fMRI task and instruction: neutral imagery with an instruction to look at the image (neutral), negative imagery with an instruction to look at the image (negative), and negative imagery with an instruction to regulate the emotional response to feel less negatively (decrease). The LASSO PCR modeling approach largely mirrors analyses from previous studies^21,24^, with adaptation to accommodate the multi-class nature of the data. The fundamentally different LDA procedure is a supervised machine learning model that utilizes linear discriminants as predictors in a multinomial logistic regression classifier without LASSO regularization. The predictive performance of these models was evaluated both via cross-validation and in a holdout dataset of 40 adults collected at a different site. Employing both models allows us to establish which method can better predict the emotional regulation task, and to compare regions that may drive this prediction.

## Results

### Predictive Performance

Figure 1 shows AUC and accuracy on six training sample sizes of the cross-validated trainingdata, intended to show the changes in modeling metrics as more training samples are provided. Assessing accuracy and AUC in cross-validation also provides evidence of the extent to which the models are overfitting to training data. The LASSO PCR model achieved an average cross-validated accuracy of 0.791 for models trained on 84% of the sample (n=69) with a corresponding AUC of 0.912. For both models, accuracy and AUC both increase as the number of training samples is increased. The LDA model slightly outperformed the LASSO PCR model at all training sample sizes in terms of both accuracy and AUC. For LDA models trained on 84% of the sample (n=69), average accuracy was 0.836 with a corresponding AUC of 0.944. Figure 1 also indicates that adding participants beyond 69 would likely produce only small increases in performance.

**Figure 1:**
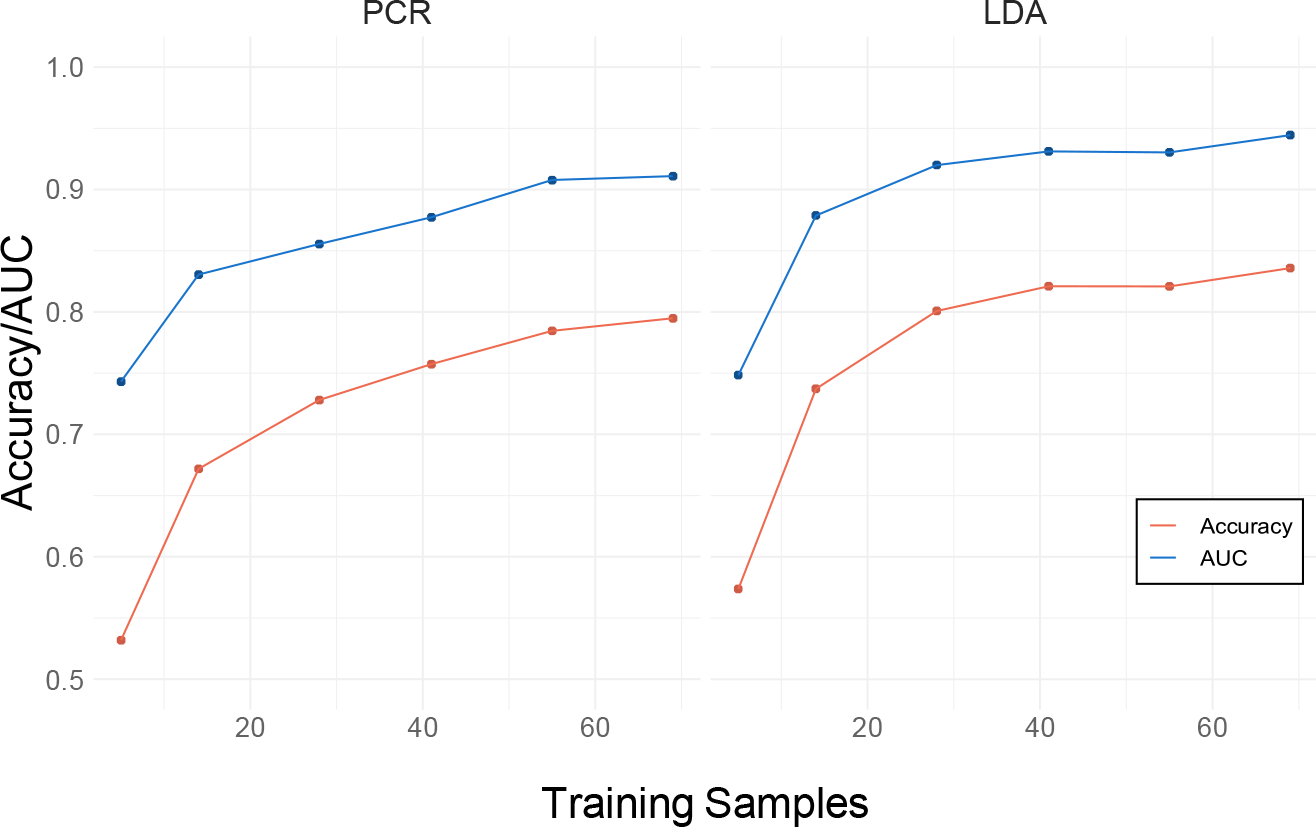
Model Cross-Validated Accuracy and AUC. Results from cross-validation are presented averaged over 20 models trained on each of six training sample sizes. The left panel displays accuracy and AUC (area under the receiver operating characteristic curve) for the LASSO PCR model while the right panel displays accuracy and AUC for the LDA model

Figure 2 displays confusion matrices of LASSO PCR and LDA classification performance in the holdout data. In the holdout data, the LASSO PCR model achieved an overall accuracy of 78.3% with a corresponding AUC of 0.916 (Table 1). Class-wise sensitivity was highest for the neutral class (0.975) and lowest for the negative class (0.625). The LDA model achieved an overall accuracy of 82.5%in the holdout data with a corresponding AUC of 0.944. No neutral class samples were misclassified, whereas class-wise sensitivity was lowest for decrease (0.675). While accuracy and AUC in the holdout data were higher for the LDA model, differences in classification performance these models in the holdout data were not statistically significant (p=0.424).

**Table 1:**
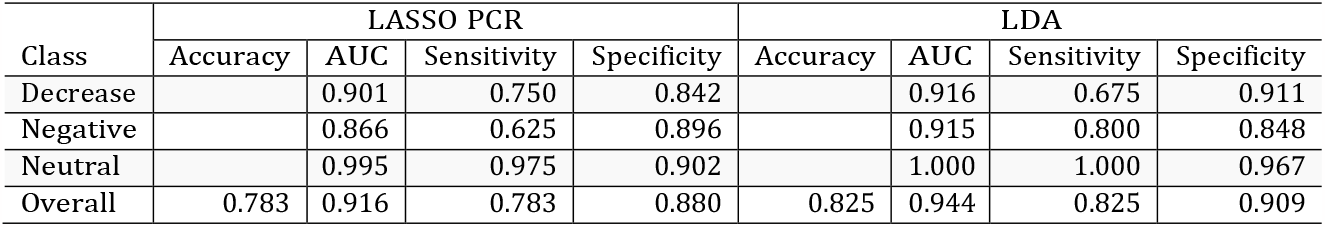
Holdout Modeling Metrics

**Figure 2:**
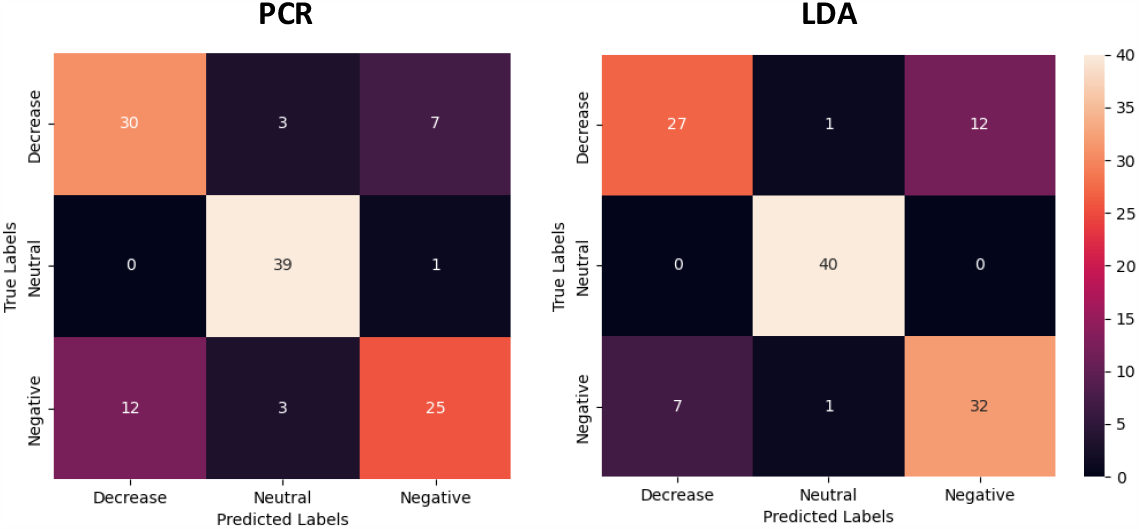
Holdout Confusion Matrices. LASSO PCR and LDA classification performances in the holdout data. Diagonals represent correctly classified samples while off diagonals contain the misclassified samples.

Figure 3 visualizes class separation for the LDA model in the training and holdout data sets. The distribution of training samples in the reduced two-dimensional feature space produced by LDA produces clear class separation, with the decrease and negative classes separated from the neutral class along the LD1 axis, while decrease is separated from negative along the LD2 axis. In order to mitigate site effects which impact the scale of linear transformations of the neural activation maps, we performed CovBat harmonization on the holdout data (Supplementary Figures 1,2). The same pattern of class separation was observed in the holdout data, with higher intraclassvariance and a greater degree of overlap between the decrease and negative classes.

**Figure 3:**
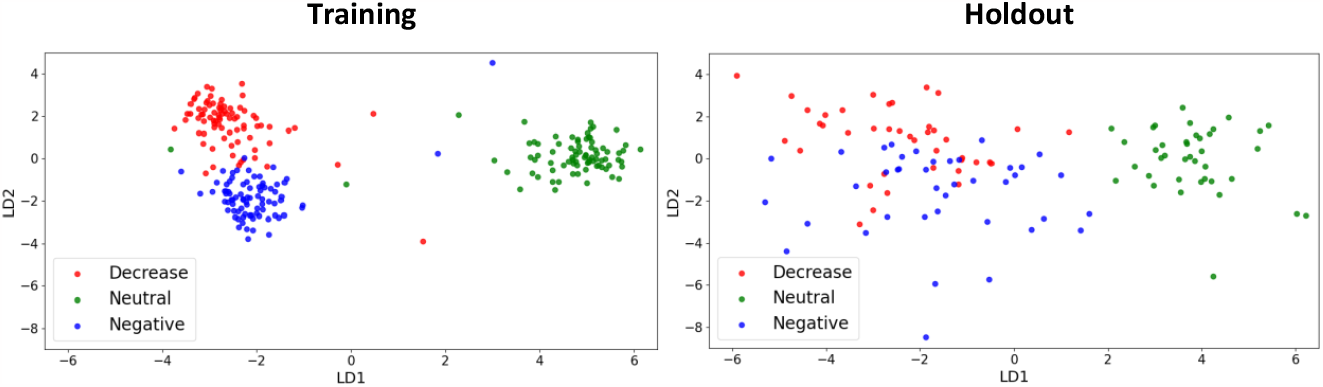
LDA Class Separation. The distribution of training samples in the reduced two-dimensional feature space produced by LDA (left panel) and the distribution of CovBat harmonized holdout samples after projection of LD loadings from training (right panel).

### Identifying the Emotional Regulation Signature

Figure 4 shows the emotional regulation signature from the LASSO PCR and LDA models, obtained via bootstrap. The voxels that are predictive of the modulation task based on the LASSO PCR model occur primarily in occipital lobe and ventrolateral PFC, while the predictive voxels derived from the LDA model occur in the insula, dorsomedial PFC, anterior lobe of cerebellum, occipital lobe, cingulate gyrus, orbitofrontal PFC, and ventrolateral PFC. Tables 2 and 3 complement Figure 4 by highlighting the brain regions of specific clusters of voxels that significantly drive class prediction for each model. These clusters of emotional regulation signature weights reveal that relatively fewer brain regions are identified as being predictively important of the emotional regulation task by the LASSO PCR model as compared to the LDA model. Both models produce clusters of weights in Brodmann Areas 6 and 40 and additionally identify prefrontal cortex as a predictively important region in the emotional regulation task.

**Table 2:**
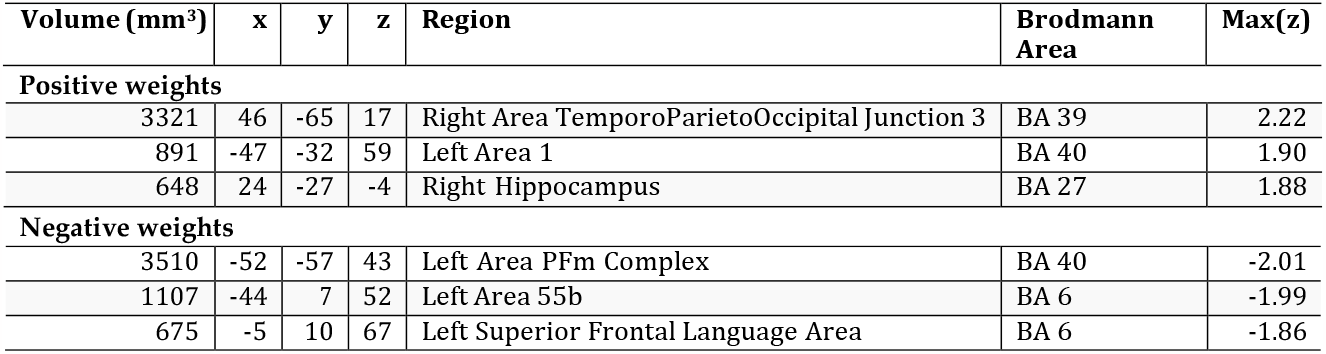
Clusters of Bootstrapped Emotional Regulation Signature Weights - PCR

**Table 3:** Clusters of Bootstrapped Emotional Regulation Signature Weights - LDA

| Volume (mm<sup>3</sup>) | x | y | z | Region | Brodmann Area | Max(z) |
| --- | --- | --- | --- | --- | --- | --- |
| **Positive weights** |  |  |  |  |  |  |
| 1890 | -49 | -61 | 47 | Left Area PFm Complex | BA 40 | 2.72 |
| 1782 | -40 | 19 | -1 | Left Middle Insular Area | BA 47 | 2.91 |
| 1296 | -53 | -37 | 3 | Left Area STSv posterior | BA 22 | 2.78 |
| 702 | -10 | 13 | 62 | Left Superior Frontal Language Area | BA 6 | 2.52 |
| 540 | 42 | -70 | -13 | Right Fusiform Face Complex | BA 19 | 2.49 |
| 513 | 45 | -49 | -22 | Right Fusiform Face Complex | BA 37 | 2.77 |
| 432 | -6 | -60 | -15 | Left Cerebellum (VI) |  | 2.82 |
| 432 | 41 | -30 | 52 | Right Area 2 | BA 3 | 2.26 |
| **Negative weights** |  |  |  |  |  |  |
| 1944 | 31 | -83 | 23 | Right Fourth Visual Area | BA 19 | -2.63 |
| 1836 | -30 | -43 | -11 | Left ParaHippocampal Area 3 | BA 37 | -3.13 |
| 1809 | 15 | -51 | 17 | Right Parieto-Occipital Sulcus Area 1 | BA 30 | -3.04 |
| 1755 | -13 | -55 | 15 | Left Parieto-Occipital Sulcus Area 1 | BA 30 | -2.83 |
| 1755 | -41 | -23 | 51 | Left Primary Sensory Cortex | BA 3 | -2.54 |
| 1512 | -49 | -23 | 15 | Left Area PFcm | BA 40 | -2.92 |
| 1026 | -34 | -84 | 28 | Left Area PGp | BA 19 | -2.50 |
| 999 | -8 | -92 | 5 | Left Primary Visual Cortex | BA 17 | -2.50 |
| 999 | 13 | -40 | 48 | Right Area 5m ventral | BA 7 | -2.72 |
| 945 | 29 | -39 | -12 | Right Ventral Visual Complex | BA 37 | -2.67 |
| 540 | 17 | -98 | -7 | Right Primary Visual Cortex | BA 17 | -2.43 |
| 513 | 49 | -33 | 30 | Right Area PFcm | BA 40 | -2.48 |
| 405 | 2 | 16 | -10 | Right Area 25 | BA 25 | -2.73 |

**Figure 4:**
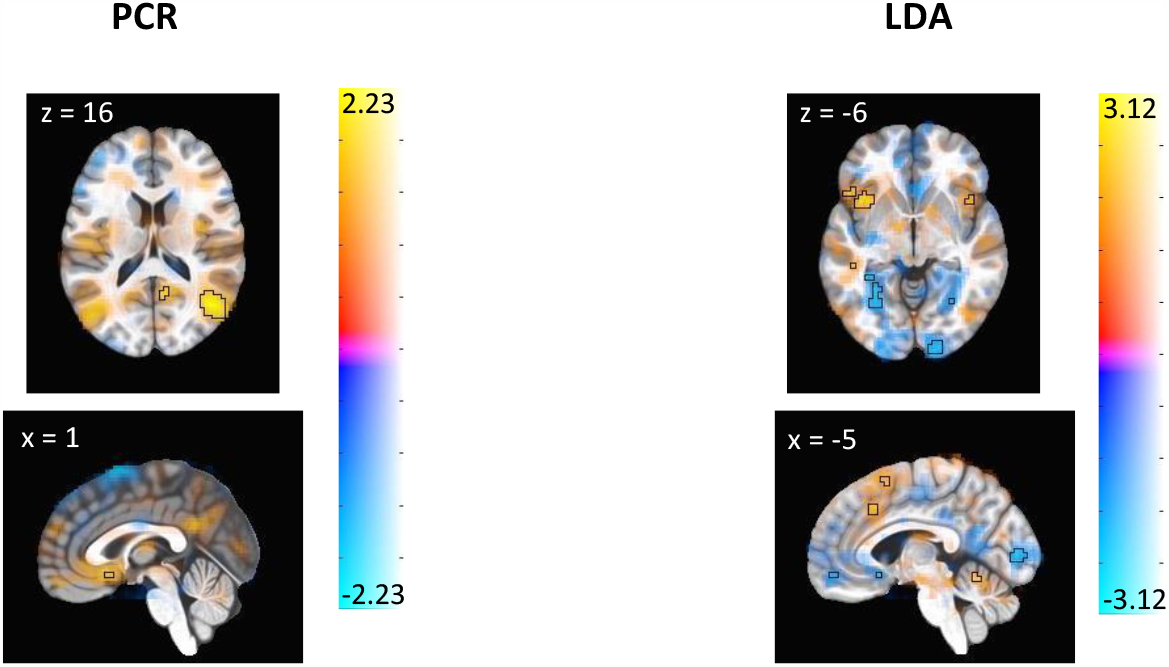
Thresholded Display of Emotional Regulation Signature. These images display brain regions which contribute most consistently to the identification of emotion regulation. Color shows the average contribution of each voxel to the prediction of emotion regulation, with warm color regions depicting positive associations and cool color regions depicting negative associations. Outlined regions display the voxels whose contributions are significant at p<0.005 uncorrected for LASSO PCR (left panel) and LDA (right panel). Note that slice location is not uniform between panels.

### Self-reported Scores of Negative Emotion

In the training sample, self-reported feelings of negativity were highest for the negative class (mean=3.502, sd=0.655) with lower reported scores for decrease (mean=2.811, sd=0.605) and neutral (mean=1.111, sd=0.196). A total of 14 subjects (17.1%) reported higher feelings of negativity during the decrease condition as compared to passive viewing of negative imagery. In the holdout sample, similar trends are observed with the highest self-reported scores of negativity observed for the negative class (mean=3.943, sd=0.926) with lower scores for decrease (mean=3.338, sd=0.926) and neutral (mean=1.141, sd=0.315). In this sample, 11 subjects (27.5%) reported higher feelings of negativity during the emotional regulation task as compared to passive viewing of negative imagery.

## Discussion

In this study, we explored the neural signatures of emotional regulation among a population of neurotypical young adults using two machine learning methods. Both methods produced accurate predictions of the fMRI task when evaluated in cross-validation and in a holdout dataset. This accuracy is noteworthy because the external stimuli are highly similar for passive viewing versus active reappraisal of negative imagery; the only difference is the internal thinking associated with cognitively changing the meaning of the negative stimuli. The predictive performance of these models indicates that despite the similarity of these stimuli, it is possible to determine when an individual engages in cognitive reappraisal using neural activation patterns.

Our signatures show contribution to the identification of emotion from some brain regions indicated in previous meta-analyses of emotion regulation, in addition to contributions from regions not identified in these reports^25,26^. For example, meta-analyses indicate that prefrontal cortex regions (ventromedial PFC, ventrolateral PFC, and others) contribute to emotion regulation, along with insular cortex and left superior frontal gyrus. We similarly identify activation from these regions as important contributors to the neural signature of emotion regulation. In concordance with prior accounts of the picture induced negative emotion signature^17^, we also identify occipital cortex regions as being important for accurate identification of emotion processing. It is plausible that the omission of visual processing regions from previous reports of emotion regulation result from biases against brain regions that have an indirect cognitive role and do not fit into theoretical neurocognitive frameworks^27^. However, it should also be considered that occipital cortex activation is likely to at least some extent specific to this visually based fMRI task, and its involvement in emotional regulation more generally is unknown.

While many previous studies have utilized LASSO PCR to identify brain regions which are associated with performing an fMRI task, we choose to additionally present the signature derived from LDA in order to provide a more comprehensive and robust representation of reappraisal. In doing so, we establish increased confidence in the involvement of brain regions identified by both models in driving the prediction of emotion regulation. Specifically, prefrontal cortex and left superior frontal gyrus are identified in the signatures produced by both models, validating prior accounts of the involvement of these regions in higher order processes^28^. While the usage of LASSO regularization reduces overfitting of the PCR model, it is important to consider the impact this has on conclusions drawn about brain regions that predict the fMRI task. We observe here that the reappraisal signature identified by LASSO PCR is sparse relative to LDA, identifying fewer involved brain regions and producing bootstrapped cluster weights with lower total volumes. While most of the information that is discarded by LASSO regularization is likely noise that is not associated with the fMRI task, there may also be relevant signal that is being missed. Consequently, some relevant brain regions which have more subtle differences in activation may not be identified in neuromarkers derived from LASSO PCR models. Although LASSO PCR is generally accepted as the gold standard method for the analysis of task-based fMRI data, these aspects suggest that comparing and contrasting results obtained from different methods may help in developing a comprehensive understanding of the underlying associations between brain regions and the fMRI task.

The modeling of a neural signature of emotion regulation meets an important criterion since it was developed on a dataset from one site but produced similar accuracy for data collected at a different site. In order to assess model performance on holdout data, it was necessary to adjust for site-specific differences in scanners and data pre-processing steps produce prior to prediction^29^. While prior work has utilized COMBAT harmonization to address site effects in functional connectivity measurements^30^, more recent work has shown that making an additional correction for feature covariance (CovBat) further reduces the impact of site effects in neuroimaging data on machine learning models^31^. In the data used inthis study, applying CovBat by class largely eliminated site-specific distributional differences in activation maps. The accuracy and AUC achieved by both models in the harmonized holdout dataset is very similar to the values these metrics achieved in cross-validation for models trained on 84% of available data. The concordance of these metrics speaks well to the generalizability of these models to unseen data, indicating that signatures which are discriminative of emotional regulation are not specific to the training sample. Notably, both models discriminate the neutral class from decrease and negative with nearly perfect accuracy, while also achieving high classification accuracy of the negative and decrease classes.

One limitation worth considering in this analysis is in the degree to which subjects were successful in performing the task of emotional modulation that is characteristic of the decrease class. That is, while self-reported feelings of negativity were lower in the decrease class on average as compared to negative, there remains a substantial subset of subjects in both the training and holdout samples who reported feeling more negatively when given the decrease-negative instruction as compared to look-negative, indicating a potential failure to regulate the emotional response. This factor could potentially reduce differences in neural signatures related to the decrease instruction and may aid in explaining the relative difficulty of correctly classifying negative and decrease samples relative to neutral, in particular given that the holdout data set contained a greater proportion of individuals for whom this factor is relevant. If individuals who report feeling more negatively during the emotional regulation task in fact have distinct signatures from those who report feeling less negatively, this effect could potentially be captured by integrating self-report information as a separate data modality during training.

The endeavor to understand the human emotional response has motivated a host of scientific papers studying brains of healthy individuals and individuals with mental health disorders^28,29^. This is not without good reason-the regulation of emotion serves a critical function in most every aspect of the human experience from the motivation of behavior to social function. This work extends our understanding of the emotional response by exploring the signatures that are unique to conscious control of emotion, while also validating previously published accounts of signatures which are characteristic of negative emotional experience. Our results further show that extensions of the LASSO principal components regression and linear discriminant analysis models to multiclass neuroimaging data produce signatures that are highly accurate at distinguishingemotional regulation tasks both in cross-validation and holdout analyses. This fMRI-based neuro-marker serves to further our understanding of the circuitry involved in cognitive reappraisal and may lend itself towards future explorations of psychopathologies and their treatment.

## Methods

Experimental procedures for the primary sample were approved by the Colorado Multiple Institutional Review Board (COMIRB) under protocol 18-1426. All participants provided written informed consent.

### Participants

The training data consists of eighty-two subjects (mean age = 20.95, SD = 1.34, 54.2% Female, 77.2% Caucasian) recruited from the Denver metro area. Participants were eligible if they were between the ages of 18 and 22 at time of study enrollment and were MRI eligible. Exclusion criteria were 1) being treated for a psychiatric disorder, 2) seeking treatment for alcohol use disorder, 3) regular tobacco use, 4) using medications that affect the hemodynamic response, 5) history of head trauma with loss of consciousness, 6) evidence of cannabis misuse (defined as >8 uses of cannabis per month or a score >11 on the Cannabis Use Disorder Identification Test-Revised), 7) >10 lifetime uses of illicit substances, 8) misuse of prescription medication in the prior year, or 9) pregnancy.

Forty subjects (mean age = 25.5, SD = 4.7, 70% Female, 87.5% Caucasian) were recruited from the Boulder, CO community for a separate study on cognitive reappraisal^30^. Subjects in this sample were given the same set of fMRI tasks, and these forty subjects were used as holdout data toassess the generalizability of the models.

### Task and Stimuli

Each subject was shown images from the International Affective Picture System (IAPS), a database containing a standardized set of images adopted widely in psychological research for the study of emotion and attention^32^. Images displayed to subjects in this study consisted of 15 neutral images and 30 negative images. The 15 neutral images were paired with a “look” instruction to the subject, indicating that the participant should simply maintain their attention on the visual stimulus. 15 negative images were paired with the same “look” instruction while the remaining 15 were paired with a “look-decrease” instruction, indicating that the participant should observe the aversive photo while attempting to consciously modulate their emotional response to feel less negatively. The instruction was shown to the subject for two seconds prior to display of the image for seven seconds. Display of the image was followed by a rest period of between one and three seconds, after which participants were given four seconds to provide a score on how negatively they felt after viewing the image on a scale from 1 (not at all negative) to 5 (very negative).

### MRI Acquisition

Images gathered for the training sample were collected on a Siemens 3.0 Tesla Skyra Magnet with a 20-channel head coil. Functional images were acquired using BOLD signal across 40 axial slices with TR = 2000-ms, TE = 30-ms, flip angle = 77, field of view = 220-mm, 40 axial 3-mm thick slices, and multiband slice acceleration factor of 2 to increase spatial resolution while maintaining temporal signal-to-noise ratio. Images were acquired in oblique orientation and high-resolution T1-weighted images were obtained for anatomical reference.

Images gathered for the holdout sample were collect on a 3.0 Tesla Siemens Magnetom Prisma system with a 32-channel head coil, with functional images acquired using BOLD signal across 56 contiguous slices with parameters TR = 460-ms, TE = 27.2-ms, flip angle = 44 °. and 3-mm thick slices^30^.

### Data Preprocessing

Data preprocessing was performed with Analysis of Functional NeuroImages (AFNI) Version 22.0.21^36^. All images were converted into AFNI-compatible file formats and echoplanar images were aligned with anatomical images. Spatial normalization was performed by nonlinearly warping the anatomical data to the Montreal Neurological Institute (MNI) standard space using AFNI SSwarper^37^. Images were skull-stripped deobliqued using the MNI152_2009_template_SSW.nii.gz template (https://afni.nimh.nih.gov/pub/dist/doc/htmldoc/template_atlas/sswarper_base.html). The final voxel resolution was 3-mm isotropic. Time points with greater than 0.3-mm Euclidean distance of framewise displacement were censored from analyses. An 8-mm kernel was used for blur, and each run was scaled to produce a mean intensity for each voxel of 100. A 7-second block model was applied to regression analysis to obtain voxel-level beta weights for each class of stimulus. Regressors of interest were the presentation of each cue (look-neutral, look-negative, decrease-negative). We included six regressors of no interest to account for motion of translation and rotation in the x, y, and z dimensions. Beta values from this regression model were then utilized as class-specific activation maps (beta maps). Images gathered for holdout analysis followeda separate preprocessing pipeline detailed in Powers et al. (2022)^34^.

### LASSO Principal Components Regression

Principal components analysis was performed on the training data, and 65 principal components were retained to reach a threshold of 90% cumulatively explained variance. These principal components were then utilized as predictors of the stimulus class in a LASSO penalized multinomial logistic regression model, where the LASSO tuning parameter *λ* was selected as the value which minimized multinomial deviance in 10-fold cross-validation. When predicting on test data, the principal component loadings from the training data were projected onto the test data.

### Linear Discriminant Analysis

Linear discriminant analysis is a method of identifying a linear combination of features to separate two or more classes of data in a manner that maximizes the ratio of between-class variance to within-class variance^38^. We utilize LDA in this context to reduce the dimensionality of the imaging features in a manner similar to the usage of PCA in the LASSO PCR model, but with the distinction that LDA dimensionality reduction explicitly considers the differences between the classes of data rather than building a reduced set of features based on feature variance, guaranteeing maximal separability^39^ LDA using two components was performed on the training data, with loadings projected onto test data for prediction. A multinomial logistic regression model was fit to the training data using the two linear discriminants as predictors. Visualizations of observations of each class were constructed to evaluate the linear separability of the classes.

### Feature Harmonization

Given that the training and holdout data were collected at different institutions with distinct pre-processing pipelines and different scanners, it was necessary to harmonize the distributions of features to mitigate site effects. To this end, CovBat^31^ was performed separately on beta maps for each class of stimulus, harmonizing the voxel-level distributions in the training and holdout data. Splitting the data by class prior to harmonization leverages the fact that we anticipate beta weight distributions at many voxels to differ by class, as do we anticipate that each class willhave a unique covariance matrix, as is evidenced by site-specific distributional differences in PC distributions by class which are mitigated after harmonization (Supplementary Figures S1, S2).

### Assessing Predictive Performance

Predictive performance was assessed on the training and test sets using measures of accuracy and area under the curve (AUC). On the training data, cross-validation was performed on each of six training sample sizes (5, 14, 28, 41, 55, 69). For cross-validation, multiclass AUC was calculated as described in Hands and Till (2001)^40^. The same method was used to calculate overall AUC for both models in the holdout dataset, while the class-wise AUCs reported in table 1 were calculated using the one-versus-rest approach. Predictive performance of the PCR and LDA models were compared in the harmonized test data using McNemar’s test.

### Voxel-wise Contributions to Class Prediction

We used the bootstrap approach described in Koban et al.^19^ to obtain voxel-level p-values and test the reliability of each voxel’s contribution to class prediction. This strategy was used for both the LASSO PCR and LDA models using the same 1000 bootstrap samples. Specifically, for each bootstrap sample a LASSO PCR and an LDA model were fit. Then, for each bootstrap sample and each class k a vector of weights the same length as the number of voxels in each image was calculated by *W*_*k*_ = *Vβk*. Here *W*_*k*_ is a vector of contribution weights for class k and *βk* is the vector of fitted multinomial logistic regression coefficients for class k. For LASSO PCA, *V* is a matrix of principle component loadings selected as nonzero by LASSO, and for LDA *V* is a 2 column matrix of linear discriminant values.

The sign of these weights were recorded across the 1000 bootstrap samples, and voxels which made consistently positive or negative contributions to the prediction of a class were identified based on a threshold of p<0.005, indicating that the voxel contributed either positively or negatively to the prediction of the decrease class at least 99.5% of the time. Voxels meeting this threshold were visualized by thresholding according to the voxel’s z-statistic from the bootstrapped distribution of contribution weights.

