## Supplementary Figures for "Neural Signatures of Emotion Regulation"

*Harmonization of Voxel Distributions*

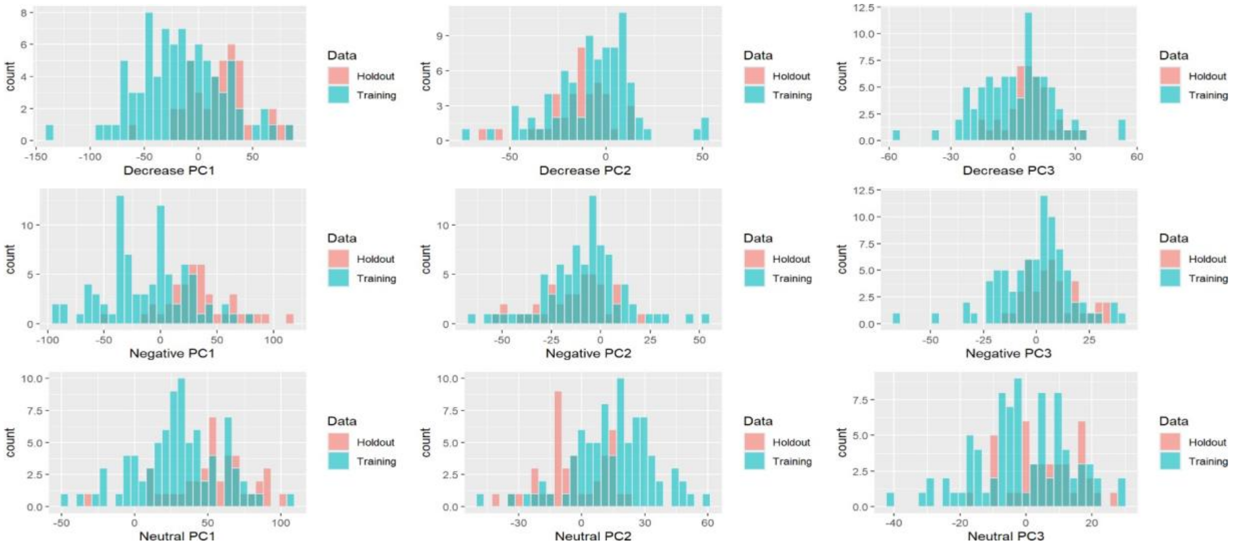


**Figure S1:** PC Distributions- Pre CovBat

Clear distributional differences are observed in the first three principal components between the training sample (blue) and holdout sample (red) prior to harmonization. The magnitude of distributional differences is modulated by class, with greater similarity observed for the negative classin PC2 and for the decrease class in PC3.

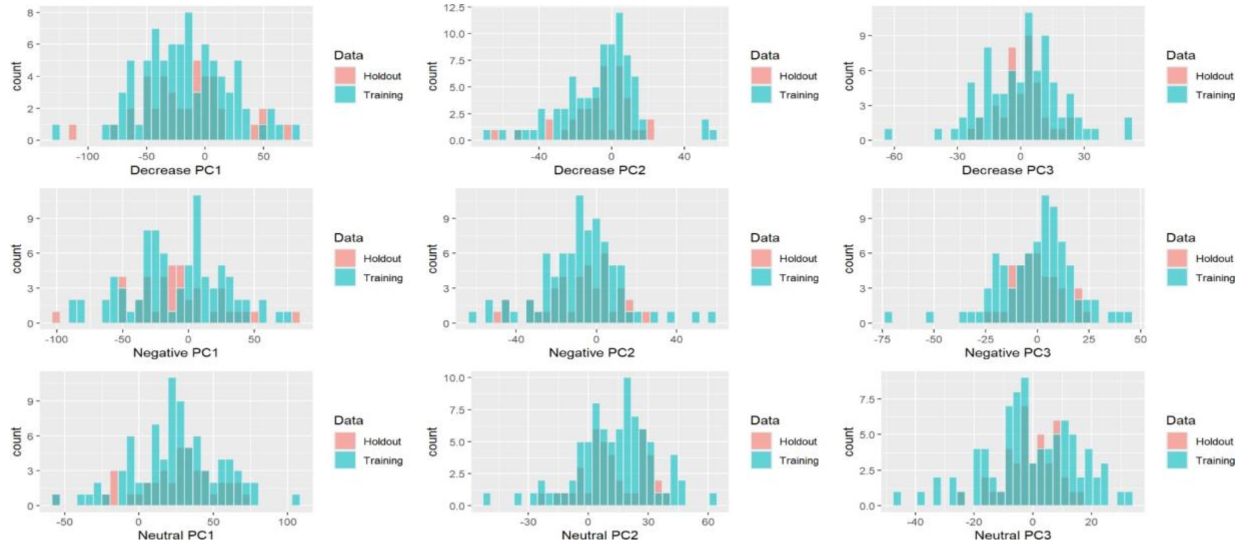


**Figure S2:** PC Distributions- Post CovBat

After CovBat harmonization, distributional differences between the training and holdout samples in the first three principal components are mitigated, with no clear class modulation effect.
